# Gene-diet interactions: dietary rescue of metabolic defects in *spen*-depleted Drosophila

**DOI:** 10.1101/770818

**Authors:** Claire M. Gillette, Kelsey E. Hazegh, Travis Nemkov, Davide Stefanoni, Angelo D’Alessandro, J. Matthew Taliaferro, Tânia Reis

## Abstract

Obesity and its co-morbidities are a growing health epidemic. Interactions between genetic background and the environment and behavior (i.e. diet) greatly influence organismal energy balance. Previously, we described obesogenic mutations in the gene *Split ends* (Spen) in *Drosophila melanogaster*, and roles for Spen in fat storage and metabolic state. In Spen-deficient storage cells lipid catabolism is impaired, accompanied by a compensatory increase in glycolytic flux and protein catabolism. Here we investigate gene-diet interactions to determine if diets supplemented with specific macronutrients can rescue metabolic dysfunction in Spen-depleted animals. We show that a high-yeast diet partially rescues adiposity and developmental defects. High sugar partially improves developmental timing as well as adult longevity. Gene-diet interactions were heavily influenced by developmental-stage-specific organismal needs: extra yeast provides benefits early in development (larval stages) but becomes detrimental in adulthood. High sugar confers benefits at both larval and adult stages, with the caveat of increased adiposity. A high-fat diet is detrimental according to all tested criteria, regardless of genotype. Whereas Spen depletion influenced phenotypic responses to supplemented diets, diet was the dominant factor in directing the whole-organism steady-state metabolome. Obesity is a complex disease of genetic, environmental, and behavioral inputs. Our results show that diet customization can ameloriate metabolic dysfunction underpinned by a genetic factor.

## INTRODUCTION

Obesity is a complex disease arising from interactions between environment, behavior, and genetics (Silventoinen *et al.* 2010; Fryar *et al.* 2016; Qasim *et al.* 2018). Metabolism of dietary nutrients into energy and biomass for survival and reproduction is a complex, highly conserved process involving three major classes of macronutrients: proteins, carbohydrates, and lipids. One major contributor to the growing obesity epidemic is Western Diet, characterized by overconsumption of calories from processed carbohydrates, fatty animal meats and byproducts, and salt, while lacking in fiber and important minerals and nutrients (Grotto and Zied 2010). Prolonged intake of excess nutrients disrupts organismal metabolic homeostasis and drives a shift from the utilization of nutrients (catabolism) to nutrient storage (anabolism).

While diet can be a major driver of fat storage, gene–environment interactions play a critical yet poorly understood role in the development of obesity and metabolic syndrome, defined as “clustering of abdominal obesity, dyslipidemia, hyperglycemia and hypertension” (Giovannucci 2007). Decades ago, the increased incidence of Diabetes Mellitus was attributed to an evolutionarily beneficial ‘thrifty’ genotype that is incompatible with the modern Western Diet and lifestyle (Neel 1962). Evidence of genetic adaptation to a diet high in polyunsaturated fatty acids can be found in the genomes of Greenlandic Inuit, in the form of genetic variants in fatty acid desaturases (Fumagalli *et al.* 2015; Chen *et al.* 2019). Accordingly, developing a comprehensive understanding of obesity and metabolic syndrome demands examination of the interplay between genetics and diet.

*Drosophila melanogaster* is an excellent model in which to study gene function in metabolism (Grönke *et al.* 2003, 2005; Baker and Thummel 2007; Leopold and Perrimon 2007; Schlegel and Stainier 2007; Guo *et al.* 2008; Beller *et al.* 2010; Reis *et al.* 2010; Hazegh *et al.* 2017; Musselman and Kuhnlein 2018). Metabolic pathways are highly conserved in Drosophila (Bernards and Hariharan 2001) and many fat regulatory genes have functional orthologues in mammals (Pospisilik *et al.* 2010; Reis *et al.* 2010). Drosophila also recapitulate the sensitivity of metabolism to genetic variation within a population while retaining the physiological plasticity to compensate, or buffer, against defects in key processes (Li *et al.* 2019; Matoo *et al.* 2019).

We have previously shown that the RNA-binding protein Split ends (Spen), a pleiotropic transcriptional regulator in Drosophila (Rebay *et al.* 2000; Lin *et al.* 2003; Hazegh *et al.* 2017), potentiates fat catabolism (Hazegh *et al.* 2017). When Spen is mutated or knocked down via RNA-interference (RNAi) in the fat body (FB; main metabolic organ of the fly) of Drosophila larvae, fat catabolism is strongly inhibited, reflected by higher levels of stored triglycerides (TAGs; Hazegh *et al.* 2017). FB-specific depletion of Spen (hereafter Spen-KD) also delays larval wandering, consistent with a failure to mobilize energy to fuel development. Impairment of fatty acid catabolism in Spen-KD larvae is accompanied by increased adiposity, decreased lipase expression, depletion of L-carnitine, and evidence of compensatory upregulation of alternative metabolic pathways, such as protein catabolism (Hazegh *et al.* 2017). Spen depletion also decreased glycogen, pointing to the use of stored carbohydrates as an auxiliary energy source in lieu of fat (Hazegh *et al.* 2017). Finally, Spen-KD larvae are short-lived under conditions of amino acid deprivation (Hazegh *et al.* 2017).

Considering the failure to utilize dietary fats and potential upregulation of protein and carbohydrate catabolism, we hypothesized that custom diets designed to suit the unique metabolic demands of Spen-KD larvae may be capable of rescuing their defects in adiposity, development, and longevity. Here we report the effects of such supplemented diets. Our results illustrate gene–diet interactions in the Spen-KD model of genetic obesity in Drosophila and support the use of dietary interventions to mitigate the influence of genetic mutations on metabolic functions and physiological outcomes. Since Spen homologs also regulate energy balance in mammals (Hazegh *et al.* 2017), our findings provide a foundation for possible dietary interventions in humans struggling with defects in metabolic pathways involving Spen.

## MATERIALS AND METHODS

### Fly Strains and Husbandry

W^1118^ (Bloomington stock number (BL) 3605), w; dcg>Gal4 (BL 7011), y^1^ sc v^1^; +; UAS-Spen RNAi (BL 33398), and y^1^ v^1^; +; UAS-w RNAi (BL 28980) were obtained from the Bloomington Drosophila Stock Center. Animals were reared at 25°C and 60% humidity unless otherwise specified and fed a modified Bloomington media (1L: yeast 15.9g, soy flour 9.2g, yellow cornmeal 67.1g, light malt extract 42.4g, agar 5.3g, light corn syrup 70mL, propionic acid 4.4mL, tegosept in ethanol 8.4mL). Medium-Yeast Diet (MYD) contained 35g yeast per liter. High-Yeast Diet (HYD) contained 70g yeast per liter. High-Fat Diet (HFD) is MYD with an additional 150g coconut oil per liter (15%). High-Sugar Diet (HSD) is MYD with an additional 180g corn syrup per liter (1M). Eggs were collected on grape plates at 25°C and 50 first-instar larvae were transferred 22-24 hours later into a vial of the specified diet.

### Density Assay

Density assays were performed as described previously with 50 larvae per sample (Reis *et al.* 2010; Hazegh and Reis 2016; Hazegh *et al.* 2017). *n*=5 for each experimental condition and genotype. Two-way ANOVA was used to calculate statistical significance (*, p<0.05) with Prism 8 software (GraphPad Software, Inc, La Jolla, CA).

### Developmental Timing

50 first-instar larvae were transferred per genotype per diet. Larvae were reared in diets upon hatching and then counted three times daily (8:00am, 12:00pm, and 4:00pm) for developmental progress. *n*=3 (50 larvae per n). Log-rank (Mantel-Cox) test and Cliff’s delta (d) were calculated to evaluate significant differences between gene-diet developmental stages, using with Prism 8 software.

### Longevity

Upon eclosion, flies were allowed 48 hours to mate, whereupon flies were separated by sex and 20 of same sex were placed into new vials. Flies were transferred every two days into vials of fresh food and scored for deaths once every day. Log-rank test was used to calculate statistical significance with Prism 8 software. n=3.

### Diet Effect Data Analysis

To determine if there was a change caused by a given gene x diet interaction, we compared the ratios of distributions across diets in one genetic background (for example, Spen-KD|MYD vs. Spen-KD|EXP) to the ratios of distributions across diets in another genetic background (for example, Ctrl-KD|MYD vs. Ctrl-KD|EXP). These two sets of ratios were then compared to each other using a Wilcoxon rank sum test. To determine the effect size of a diet’s effect on a given genotype, Cliff’s delta was calculated on the two sets of ratios. Values for Cliff’s delta range between −1 and 1, and larger value absolute values indicate stronger effects. Statistical tests and Cliff’s delta calculations were performed using a custom Python script.

### Metabolomics

Briefly, individual larvae (10 per sample, n=3 samples per genotype) were suspended in 1 ml of methanol/acetonitrile/water (5:3:2, v/v) pre-chilled to −20°C and vortexed continuously for 30 min at 4°C. Insoluble material was removed by centrifugation at 10,000xg for 10 min at 4°C and supernatants were isolated for metabolomics analysis by UHPLC-MS. Analyses were performed as previously described (Nemkov *et al.* 2017, 2019) using a Vanquish UHPLC system coupled to a Q Exactive mass spectrometer (Thermo Fisher Scientific, San Jose, CA, USA. Orthogonal partial least squares discriminant analysis (OPLS-DA) was used to compare differences within both the genotype and between genotypes on the same diet. Graphs, heat maps and statistical analyses (t-Test) were performed with GraphPad Prism 8.0.

Flies are available upon request. All data necessary for confirming the conclusions of the article are present within the article, figures, tables and supplemental data.

## RESULTS

### High-sugar and high-yeast diets dampen the effect of fat body-specific Spen-depletion on fat storage

Spen-KD causes an increased adiposity phenotype on the control diet (Medium-Yeast Diet; MYD) compared to the White-RNAi control (CTRL-KD; Figure 1A, Hazegh *et al.* 2017). To test if macronutrient supplementation can rescue the increased fat storage in Spen-KD larvae, the standard MYD was supplemented with excess yeast to make a High-Yeast Diet (HYD) as used in previous studies on diet composition and lifespan (Lee *et al.* 2008; Skorupa *et al.* 2008). Yeast is the main source of protein in standard fly diets, though it also provides other nutrients, such as vitamins and carbohydrates. Supplementation with 1M sugar made a High-Sugar Diet (HSD) as used in previous studies examining Drosophila obesity and insulin resistance (Musselman *et al.* 2011). We used a High-Fat Diet (HFD) supplemented with coconut oil, as used in previous obesity pathophysiology research in Drosophila (Birse *et al.* 2010). Larvae were transferred from grape plates into these diets as first-instar larvae, at 22-24 hours after egg deposition (AED), and collected at the third-instar wandering stage to measure buoyancy (Figure 1). There are sex-specific differences in adiposity, however the Spen-KD effected both sexes similarly (Hazegh *et al.* 2017), therefore we analyzed mixed male/female larvae in the buoyancy assays. For all buoyancy assays, progeny from crosses of UAS-Spen RNAi, UAS-W RNAi, and dcg>GAL4 to w1118 strains were used as genetic background controls (Supplemental Figure 1, Hazegh *et al.* 2017).

**Figure 1:**
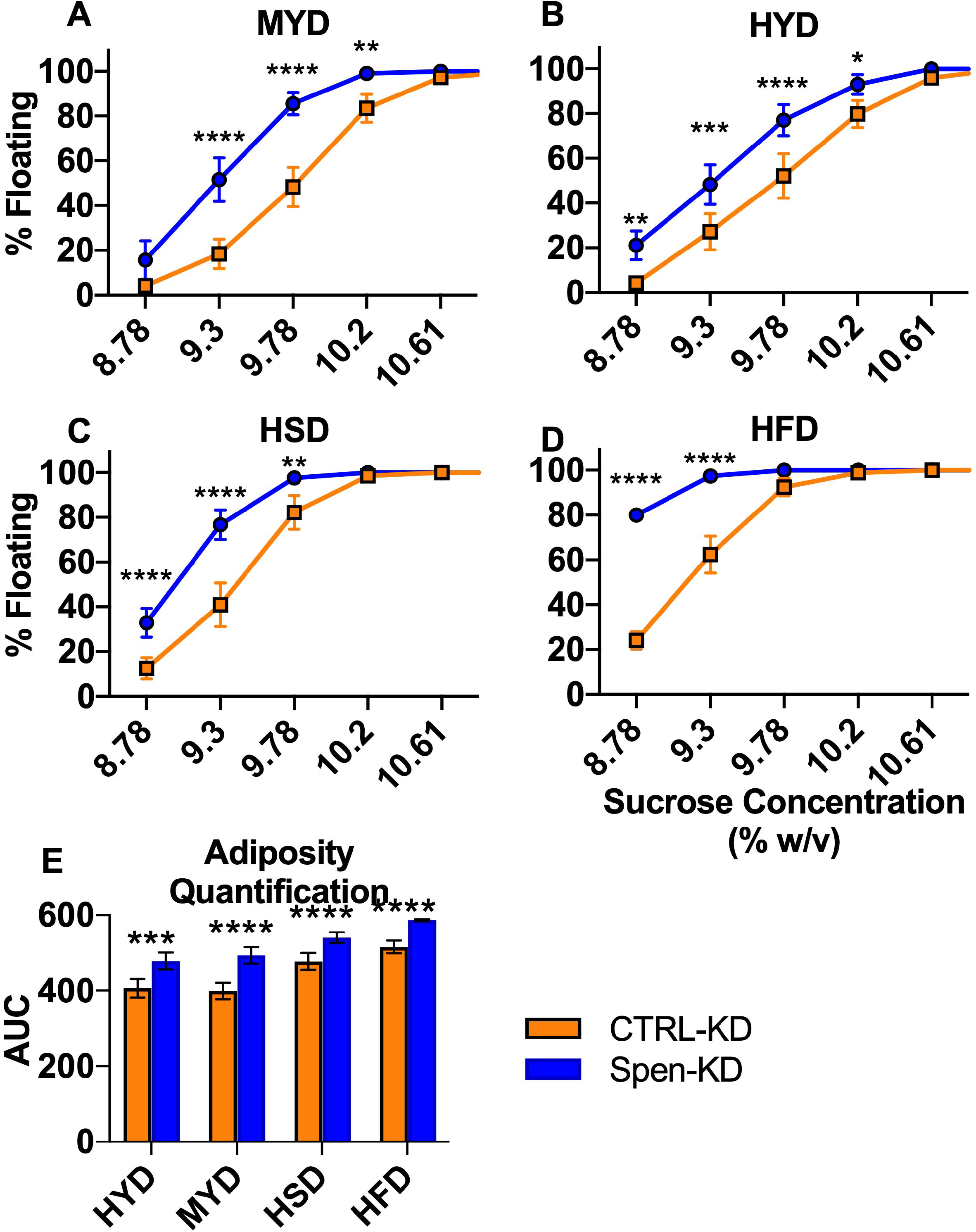
Dietary intervention affects the adiposity phenotype of Spen-KD. Buoyancy assays were performed on third-instar wandering larvae; 50 larvae per genotype per experimental replicate, n=5. Error bars represent SEM. A) The percent of floating larvae at different sucrose densities (percent weight/volume) of FB-specific Spen KD (Spen-KD, blue) compared to control KD (CTRL-KD, orange) reared on the control Medium-Yeast Diet (MYD). B) As in Figure 1A, Spen-KD buoyancy compared to CTRL-KD across a sucrose gradient with larvae reared on High-Yeast Diet (HYD), C) reared on High-Sugar Diet (HSD), and D) reared on High-Fat Diet (HFD). Two-way ANOVA, *p<0.05, **p<0.01, ***p<0.001, ****P<0.0001. E) Quantification of the buoyancy curves by integrating the area under the curve (AUC) represented as arbitrary units (A.U.). Comparison by two-way ANOVA (***p<0.001, ****p<0.0001).

We compared the buoyancy of the larvae on different diets by integrating the area under the curves (AUC) generated by plots of percent floating larvae versus sucrose concentration and comparing the differences between areas (Figure 1E, Table 1). Spen-KD was significantly fatter than CTRL-KD on all diets (two-way ANOVA, p<0.05; Figure 1A-D). MYD drove the largest buoyancy difference between Spen-KD and CTRL-KD (94.7 AU; Figure 1A, Table 1). Both Spen-KD and CTRL-KD had the largest AUC on HFD, indicating the HFD drives the largest increase in adiposity regardless of genotype (Figure 1D, Table 1). The smallest difference between Spen-KD and CTRL-KD was observed on the HSD (Figure 1C, Table 1). While HSD increased the adiposity of Spen-KD and CTRL-KD when compared to MYD matched controls, this diet decreased genotypical adiposity differences (Table 1).

**Table 1:** Quantification of the area under the curve (AUC) for buoyancy assays of Spen-KD and CTRL-KD on diets. A.U., Arbitary Units. High-yeast diet (HYD), Medium-yeast diet (MYD), High-sugar diet (HSD), High-fat diet (HFD). Two-way ANOVA analysis, ***p<0.001, ****p<0.0001. It is important to note that HFD increased adiposity outside of the linear range.

| Diet | CTRL-KD<br>AUC A.U. | Spen-KD<br>AUC A.U. | Area Difference<br>Spen-KD - CTRL-KD | p-value |
| --- | --- | --- | --- | --- |
| MYD | 399.3 | 494 | 94.70 ± 11.75 | <0.0001 |
| HYD | 407 | 479 | 72.00 ± 12.71 | <0.001 |
| HSD | 478 | 540.9 | 62.90 ± 10.01 | <0.0001 |
| HFD | 515.9 | 587.4 | 71.50 ± 7.558 | <0.0001 |

### High-sugar and high-yeast diets partially rescue previously characterized developmental delays in Spen KD

Larvae in which Spen is depleted in the FB are delayed in development (Hazegh *et al.* 2017). To test how diet supplementation affects development, Spen-KD and CTRL-KD larvae were reared in HYD, HFD, or HSD from first-instar larvae to eclosion. Vials were assessed three times daily for numbers of wandering larvae, pupal cases, and eclosed flies and compared by two-way ANOVA, Cliff’s delta, Log-rank (Mantel-Cox), and Chi-square analysis to examine gene-diet interaction.

Timing of third-instar wandering stage was more affected by the genotype than by diet composition (Figure 2). The median wandering time of Spen-KD|MYD larvae was 152.7 hours AED, a significant 24.7 hour delay from the CTRL-KD|MYD (wandering median of 128 hours AED; two-way ANOVA, p<0.0001; Figure 2A). The delay observed between the Spen-KD|EXP and CTRL-KD|EXP diet-matched controls was strongly conserved in the different diets; Spen-KD|HYD was delayed 22.8 hours (two-way ANOVA p<0.0001; Figure 2A), Spen-KD|HSD was delayed 23.5 hours (two-way ANOVA, p= p<0.0001; Figure 2B), and Spen-KD|HFD was delayed 28.9 hours (two-way ANOVA, p= p<0.0001; Figure 2C).

**Figure 2:**
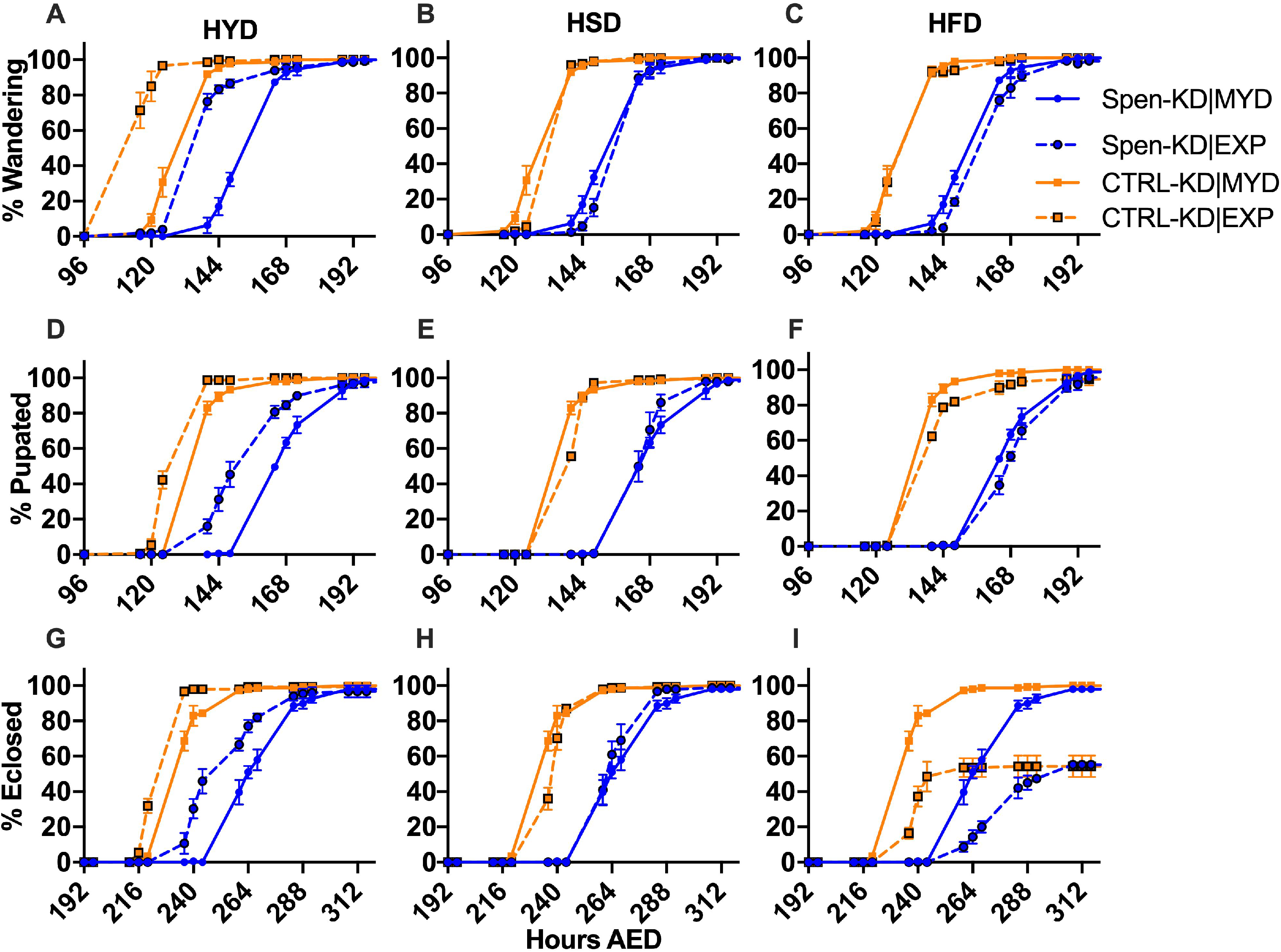
High-Yeast Diet improves developmental delays in Spen-KD larvae. Eggs were collected using the standard egg collection protocol. At 22-24 hours after egg deposit (AED) 50 larvae per genotype, per biological replicate were transferred into either into the control Medium-Yeast Diet (MYD), or an experimental diet: High-Yeast Diet (HYD), High-Sugar Diet (HSD), or High-Fat Diet (HFD). Vials were monitored at 8:00am, 12:00pm and 4:00pm for developmental progress. A-C) Spen-KD|MYD (solid blue) and CTRL-KD|MYD (solid orange) wandering percentage at each time point in hours AED compared to Spen-KD and CTRL-KD larvae reared on the experimental A) HYD, B) HSD, and C) HFD. A-C) Spen-KD|MYD (solid blue) and CTRL-KD|MYD (solid orange) pupariation percentage at each time point in hours AED compared to Spen-KD and CTRL-KD larvae reared on the experimental D) HYD, E) HSD, and F) HFD. G-I) Spen-KD|MYD (solid blue) and CTRL-KD|MYD (solid orange) eclosion percentage at each time point in hours AED compared to Spen-KD and CTRL-KD larvae reared on the experimental, G) HYD, H) HSD, and I) HFD.

To determine if the Spen-KD genotype is more responsive to a diet than CTRL-KD, we looked at effect size, a quantitative measure of the magnitude of a developmental time difference. A larger absolute value of an effect size indicates a stronger effect between the two variables. We used Cliff’s delta (d) to measure how often the values in one distribution (ratio of Spen-KD|EXP /Spen-KD|MYD) were larger than the values in a second distribution (ratio of CTRL-KD|EXP /CTRL-KD|MYD). Importantly, this does not require assumptions about how the data are distributed. d values of 0.11, 0.28, and 0.43 denote small, medium, and large effect size respectively (Vargha and Delaney 2000).

Spen-KD|HSD wandering development was not statistically different than Spen-KD|MYD, however CTRL-KD|HSD was delayed (4 hours from control, Chi square=14.95, p=0.0001; Table 2). HSD had the smallest effect size (d=0.05) between Spen-KD and CTRL-KD, indicating that a carbohydrate-enriched diet does not have as significant effect on wandering in either genotype (Table 2). Spen-KD|HFD was delayed in wandering compared to Spen-KD|MYD (5.2 hours from control, Chi square=8.459, p=0.0036), however CTRL-KD wandering was not significantly affected (Table 2). In accordance with these results, the effect size (d=-0.15) suggests that in terms of developmental timing Spen-KD is more negatively affected by HFD than CTRL-KD at the wandering stage (Table 2). Both genotypes wandered earlier when reared on the HYD compared to MYD, however CTRL-KD|HYD (15.5 hours earlier than control diet, Chi square=208.3, p<0.0001; Table 2) was more responsive to increased protein than the Spen-KD|HYD (17.4 hours earlier than control diet, Chi square= 83.47, p<0.0001; Table 2). This conclusion is supported by the effect ratio (d=-0.25), which means that both genotypes were positively affected by HYD, however CTRL-KD was more strongly affected than Spen-KD (Table 2).

**Table 2:** Developmental timing of Spen-KD and CTRL-KD genotypes on diets. Chi square from Log-rank (Mantel-Cox) test, *p<0.05, **p<0.01, ***p<0.001, ****p<0.0001, ns not significant. ^#^:Gehan-Breslow-Wilcoxon survival curve analysis, which weights earlier events more heavily due to inconsistent hazard ratios and one group with consistently higher risk. Cliff’s delta measuring how often the Spen-KD median development ratio (diet effect on the Spen-KD genotype) is higher than the values in the distribution of CTRL-KD median development ratio on the same diet. Medium-Yeast Diet (MYD), High-Yeast Diet (HYD), High-Sugar Diet (HSD), High-Fat Diet (HFD). Mean hour After Egg Deposit (AED). NA, Not applicable.

|  | Diet | CTRL-KD<br>(median<br>hours<br>AED) | Spen-KD<br>(median<br>hours<br>AED) | Spen-KD/<br>CTRL-KD<br>Delay<br>(hours) | CTRL-KD<br>(MYD\|EXP)<br>Chi<br>square | p-value | Spen-KD<br>(MYD\|EXP)<br>Chi square | p-value | CTRL-KD<br>Ratio<br>(Median<br>Log2) | Spen-KD<br>Ratio<br>(Median<br>Log2) | Ratio<br>Difference<br>p-value | Cliff's<br>delta<br>(d) |
| --- | --- | --- | --- | --- | --- | --- | --- | --- | --- | --- | --- | --- |
| Wandering | MYD | 128 | 152.7 | 24.7 | NA | NA | NA | NA | NA | NA | NA | NA |
|  | HYD | 112.5 | 135.3 | 22.8 | 208.3 | <0.0001 | 83.47 | <0.0001 | 0.263 | 0.263 | <0.0001 | -0.25 |
|  | HSD | 132 | 155.5 | 23.5 | 14.95 | <0.0001 | 2.910 | ns | 0 | 0 | 0.0001 | 0.05 |
|  | HFD | 128.6 | 157.5 | 28.9 | 0.7895 | <0.0001 | 8.459 | 0.0036 | 0 | 0 | ns | -0.15 |
| Pupariation | MYD | 135 | 164.6 | 29.6 | NA | NA | NA | NA | NA | NA | NA | NA |
|  | HYD | 124.5 | 150.6 | 26.1 | 61.59 | <0.0001 | 65.06 | <0.0001 | 0.041 | 0.148 | <0.0001 | 0.16 |
|  | HSD | 139.5 | 163.9 | 24.4 | 5.153 | <0.0001 | 5.875 | 0.0154 | 0 | 0 | 0.0232 | 0.22 |
|  | HFD | 137.1 | 167.5 | 30.4 | 4.315 | 0.09454 | 2.835 | ns | 0 | 0 | 0.0378 | 0.01 |
| Eclosion | MYD | 232.8 | 264 | 31.2 | NA | NA | NA | NA | NA | NA | NA | NA |
|  | HYD | 221.4 | 249.8 | 28.4 | 70.92 | <0.0001 | 52.44 | <0.0001 | 0 | 0.114 | <0.0001 | 0.44 |
|  | HSD | 237.6 | 262 | 24.4 | 1.666 | <0.0001 | 7.247 | 0.0071 | -0.024 | 0 | ns | 0.27 |
|  | HFD | 238.1 | 274 | 35.9 | 14.69<sup>#</sup> | <0.001 | 14.97<sup>#</sup> | 0.0001 | -0.024 | -0.022 | 0.0001<sup>#</sup> | -0.07 |

Spen-KD|MYD pupariation occurred at a median 164.6 hours AED, a 29.6 hour delay from CTRL-KD|MYD pupation (two-way ANOVA, p<0.0001; Figure 2D, Table 2). As observed in wandering, the delay in pupariation between the Spen-KD|EXP and CTRL-KD|EXP diet-matched controls was strongly conserved amongst the experimental diets; Spen-KD|HYD was delayed 26.1 hours (two-way ANOVA, p<0.0001; Figure 2D, Table 2), Spen-KD|HSD was delayed 24.4 hours (two-way ANOVA, p<0.0001; Figure 2E, Table 2), and Spen-KD|HFD was delayed 30.4 hours (two-way ANOVA, p<0.0001; Figure 2F, Table 2). We next examined the sensitivity of the Spen-KD genotype to macronutrient content and asked if a partial phenotypic rescue is possible through dietary supplementation. Spen-KD|HSD pupariated earlier than Spen-KD|MYD (0.7 hours earlier than control, Chi square=5.875 p=0.0154; Table 2). Conversely, CTRL-KD|HSD was delayed compared to CTRL-KD|MYD (4.5 hours from control, Chi square=5.153, p=0.0232; Table 2). When comparing the effect of diet between genotypes, we saw that HSD had a larger effect on Spen-KD than CTRL-KD (d=0.22; Table 2). There was no significant difference in pupariation timing between Spen-KD|HFD and Spen-KD|MYD, however CTRL-KD|HFD was delayed compared to CTRL-KD|MYD (2.1 hours from control, Chi square=4.315, p=0.0378; Table 2). The HFD had a very small effect size, with no significant difference in Spen-KD but negatively affecting CTRL-KD (d=0.01). Spen-KD|HYD pupariated earlier than Spen-KD|MYD (14 hours earlier than control, Chi square=65.06, p<0.0001). The CTRL-KD|HYD also pupariated earlier than CTRL-KD|MYD (9.5 hours earlier than control, Chi square=61.59, p<0.0001; Table 2). While both genotypes responded positively to HYD, HYD had a stronger effect on CTRL-KD (d=0.16).

Spen-KD|MYD eclosed a median 264 hours AED, a 31.2-hour delay relative to CTRL-KD|MYD (two-way ANOVA, p<0.0001; Figure 2G, Table 2). Eclosion appears to be more sensitive to macronutrient content than wandering or pupariation, which could be a cumulative effect from delays in earlier development. Spen-KD|HSD eclosed an average of 2 hours earlier than Spen-KD|MYD (Chi square=7.247, p=0.0071; Figure 2H, Table 2). CTRL-KD|HSD was delayed compared to CTRL-KD|MYD (4.8 hours delayed, Chi square=1.666, p<0.0001; Figure 2H; Table 2). HSD affected Spen-KD more than CTRL-KD (d=0.27). Due to the significant lethality of the HFD at eclosion and the severity of the delay, we analyzed the HFD development timing using the Gehan-Breslow-Wilcoxon test, which weights earlier events more heavily. This test is appropriate for a data set with inconsistent hazard ratios and one group with consistently higher risk (e.g. Spen-KD early death). Both Spen-KD|HFD (10 hours delayed, Chi square=14.97, p=0.0001; Figure 2I, Table 2) and CTRL-KD|HFD (5.3 hours delayed, Chi square=14.69, p=0.0001; Figure 2I, Table 2) eclosed later than the MYD diet-matched controls. The Spen-KD genotype was more adversely affected by HFD than CTRL-KD (d=-0.07). Both Spen-KD|HYD (14.2 hours earlier, Chi square=52.44, p<0.0001; Figure 2I, Table 2) and CTRL-KD|HYD (11.4 hours earlier, Chi square=70.92, p<0.0001) eclosed earlier than their MYD controls (Figure 2G; Table 2). The effect of the HYD was larger on the Spen-KD genotype compared to the CTRL-KD (d=0.44), suggesting Spen-KD is more responsive to HYD than CTRL-KD. Despite this, Spen-KD genotype was significantly delayed in all diets. These findings point to a defect in Spen-KD animals in maintaining normal developmental timing when challenged with suboptimal diets.

### High-sugar diet partially rescues the early death of Spen-KD flies

Just as defects in metabolism can lead to delays in development, so can they lead to longevity defects (Lushchak *et al.* 2012; Kitada *et al.* 2019). Since Spen-KD flies displayed defects in development during the larval stage in conjunction with the larval fat phenotype, we asked whether there might be a long-term effect on the lifespan of adult flies. Indeed, the CTRL-KD|MYD mated females lived 84% longer than Spen-KD diet-sex-matched flies (median 68 days, compared to 37 days; Figure 3A; Table 3). Inversely, Spen-KD|MYD males lived 3% longer than CTRL-KD diet- and sex-matched flies (median 50 days, compared to 48.5 days; Figure 3D; Table 3). On MYD, CTRL-KD mated females lived longer than CTRL-KD males (median 68 days, compared to 48.5 days; Table 3), whereas Spen-KD mated females had shorter lifespans than Spen-KD males (median 37 days, compared to 50 days; Table 3).

**Figure 3:**
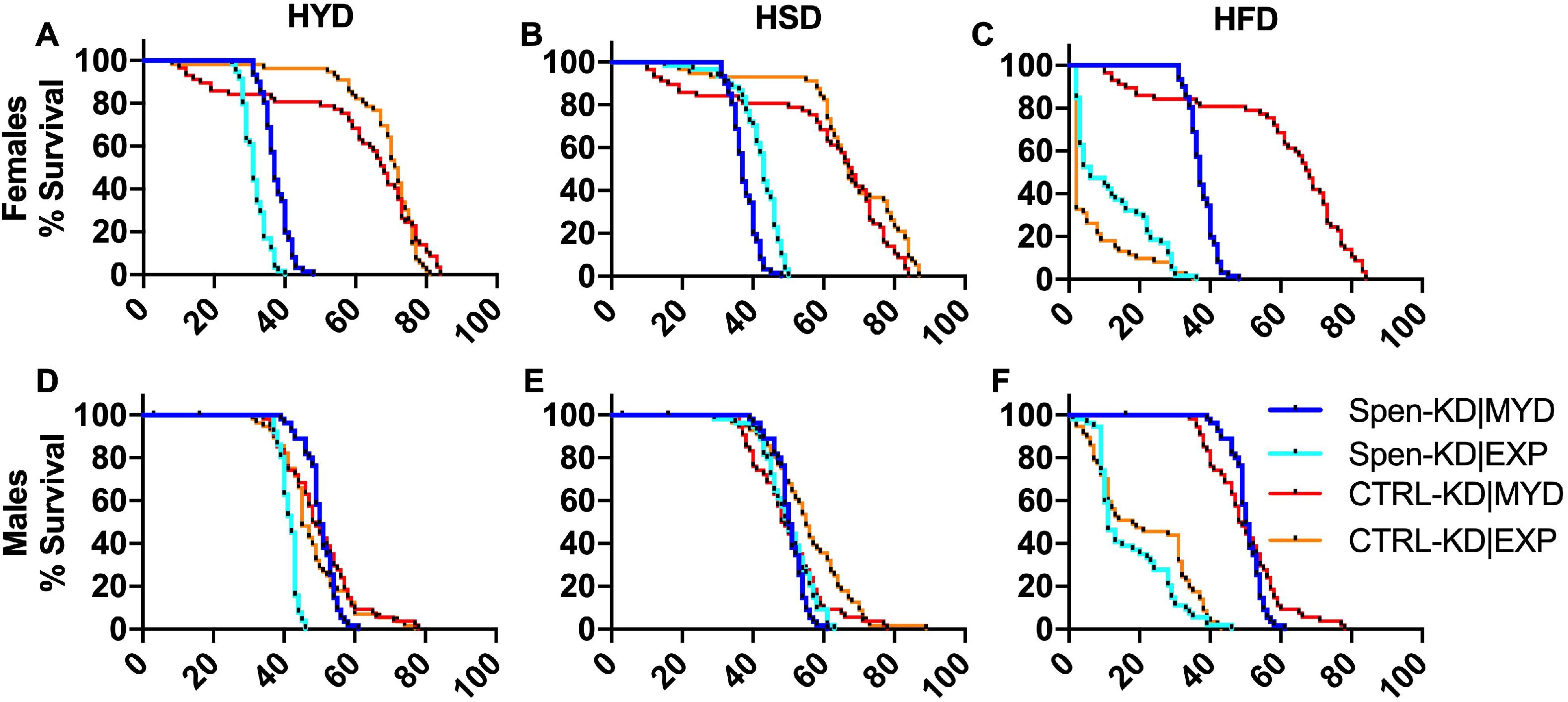
Macronutrient composition differentialy affects lifespan. Adults were allowed to eclose and mate for 48 hours before being separated by sex and transferred to either Medium-Yeast Diet (MYD) or an experimental diet (EXP): High-Yeast Diet (HYD), High-Sugar Diet (HSD), and High-Fat Diet (HFD). Vials were assessed daily for death events to generate survival curves. A-C) Mated female percent survival (days) of Spen-KD|MYD and CTRL-KD|MYD compared to A) HYD, B) HSD, and C) HFD. D-F) Male percent survival (days) of Spen-KD|MYD and CTRL-KD|MYD compared to D) HYD, E) HSD, and F) HFD.

**Table 3:** Macronutrient content of diet modulates the longevity of male and female flies. Median days elapsed to death for adult flies of the specified genotype reared in a Medium-Yeast Diet (MYD), High-Yeast Diet (HYD), High-Sugar Diet (HSD), High-Fat Diet (HFD). Chi square from Log-rank (Mantel-Cox) test, *p<0.05, **p<0.01, ***p<0.001, ****p<0.0001. Cliff’s delta (d) shows effect size of diet on genotype. NA, Not Applicable.

|  | Diet | CTRL-KD<br>Median Survival<br>(Days) | Spen-KD<br>Median Survival<br>(Days) | %<br>(Spen-KD/CTRL-KD) | CTRL-KD<br>(MYD\|EXP)<br>Chi-Square | p-value | Spen-KD<br>(MYD\|EXP)<br>Chi-Square | p-value | Cliffs<br>delta<br>(d) |
| --- | --- | --- | --- | --- | --- | --- | --- | --- | --- |
| Female | MYD | 68 | 37 | 84 | NA | NA | NA | NA | NA |
|  | HYD | 71.5 | 31 | 131 | 0.02097 | 0.885 | 56.81 | <0.0001 | 0.62 |
|  | HSD | 67 | 43 | 56 | 4.349 | 0.037 | 38.35 | <0.0001 | -0.12 |
|  | HFD | 2 | 6 | -67 | 120.5 | <0.0001 | 136.7 | <0.0001 | -0.54 |
| Male | MYD | 48.5 | 50 | -3 | NA | NA | NA | NA | NA |
|  | HYD | 45 | 42 | 7 | 0.9183 | 0.338 | 90.88 | <0.0001 | 0.37 |
|  | HSD | 55 | 50 | 10 | 6.227 | 0.013 | 1.188 | 0.276 | 0.27 |
|  | HFD | 18 | 11 | 64 | 105.0 | <0.0001 | 129.0 | <0.0001 | 0.12 |

Because HYD rescued the fat storage phenotype, and HYD and HSD were able to partially rescue various developmental delays, we predicted that altering the diet of adult Spen-KD flies might similarly improve lifespan. To test this prediction, Spen-KD and CTRL-KD larvae were reared in MYD, allowed to eclose and mate for 48 hours, separated by sex, transferred onto MYD, HYD, HFD, or HSD, and analyzed daily for survival. We observed significant lifespan differences between Spen-KD and CTRL-KD flies between diets, as well as sex- and genotype-specific responses to diets. Chi-square values for experimental diets were calculated using the MYD matched control as the predicted value.

CTRL-KD mated females had increased longevity compared to CTRL-KD males on MYD, HYD, and HSD, but not HFD (Figure 3A-C; Table 3). This trend was the opposite for Spen-KD; Spen-KD mated females had decreased longevity compared to Spen-KD males on all diets tested (Figure 3; Table 3). CTRL-KD males significantly outlived Spen-KD males on all diets except for MYD, which is statistically insignificant (Figure 3D-F; Table 3).

Spen-KD|HYD mated female lifespan was decreased 16% compared to Spen-KD|MYD controls (Chi square=56.81, p<0.0001; Figure 3A, Table 3). CTRL-KD|HYD mated female longevity was not significantly different than CTRL-KD|MYD (Chi square=0.021, p=0.885; Figure 3A, Table 3). Comparing diet-matched conditions the CTRL-KD|HYD mated females lived 130% longer than Spen-KD|HYD mated females (d=0.62; Figure 3A, Table 3). HYD had a large deleterious effect on Spen-KD mated females and no effect on CTRL-KD mated females (Figure 3A, Table 3).

CTRL-KD|HYD males lived 7% longer than Spen-KD|HYD males (two-way ANOVA, p<0.0001; Figure 3D). Spen-KD|HYD male longevity was decreased 16% compared to Spen-KD|MYD controls (Chi square=90.88, p<0.0001; Figure 3D, Table 3). CTRL-KD|HYD male longevity is not significantly different than CTRL-KD|MYD (Chi square=0.918, p=0.338; Figure 3D, Table 3). Comparing diet-matched conditions, HYD was more deleterious to Spen-KD males than to CTRL-KD (d=0.37).

CTRL-KD|HSD mated females lived 56% longer than Spen-KD|HSD mated females (Chi square=105.4 value, p<0.0001; Figure 3B, Table 3). Spen-KD|HSD mated female longevity increased by 16% compared to Spen-KD|MYD controls (Chi square=38.35, p<0.0001; Figure 3B, Table 3), and CTRL-KD|HSD mated female longevity increased 1.5% compared to CTRL-KD|MYD (Chi square=4.349, p=0.037; Figure 3B, Table 3). Comparing diet-matched conditions, the CTRL-KD|HSD mated females lived 56% longer than Spen-KD|HSD mated females (two-way ANOVA, p<0.0001, d=-0.12; Figure 3B, Table 3). Spen-KD mated females were more positively affected by HSD than CTRL-KD mated females (Figure 3B, Table 3).

We observed that HSD had no effect on the longevity of Spen-KD males, however HSD was beneficial to CTRL-KD males, which lived 13% longer than CTRL-KD|MYD (Chi square=6.227, p=0.013; Figure 3E, Table 3). CTRL-KD|HSD males lived 10% longer than Spen-KD|HSD males (d=0.27; two-way ANOVA, p<0.0001; Figure 3E, Table 3).

HFD was deleterious to the longevity of mated females, regardless of genotype. Spen-KD|HFD mated female lifespan was decreased 84% compared to Spen-KD|MYD controls (Chi square=136.7, p<0.0001; Figure 3C, Table 3), and CTRL-KD|HFD mated female lifespan was decreased 97% compared to CTRL-KD|HFD (Chi square=120.5, p<0.0001; Figure 3C, Table 3). Comparing diet-matched conditions, the Spen-KD|HFD mated females lived 67% longer than CTRL-KD|HFD mated females (d=-0.54; Figure 3C, Table 3). Interestingly, HFD was more deleterious to CTRL-KD mated females than to Spen-KD mated females (Figure 3C, Table 3).

Spen-KD|HFD male lifespan was decreased 78% compared to Spen-KD|MYD controls (Chi square=129, p<0.0001; Figure 3F, Table 3), and CTRL-KD|HFD male lifespan was decreased 63% compared to CTRL-KD|MYD (Chi square=105, p<0.0001; Figure 3C, Table 3). Comparing diet-matched conditions, the CTRL-KD|HFD males lived 64% longer than Spen-KD|HFD males (d=0.12; Figure 3F, Table 3). HFD was more deleterious to Spen-KD males than to CTRL-KD males. These findings illustrate that the effects of diet on longevity vary greatly by diet, genotype and sex. Among these effects, the apparent benefits of the HSD to Spen-KD females are consistent with our hypothesis that specific diets may ameliorate the adverse consequences of Spen depletion.

### Dietary effects on metabolism measured by steady-state metabolimic profiling

We previously showed major alterations in the metabolomic profiles of Spen-KD larvae compared to CTRL-KD on the control MYD (Hazegh *et al.* 2017). These results suggested an increase in protein catabolism and glycogen utilization and drove us to investigate dietary intervention as a treatment for the Spen-KD obesity phenotype.

L-carnitine is necessary for the β-oxidation of fatty acids within the mitochondrial matrix to feed the tricarboxylic acid cycle (TCA) (Figure 4E). L-carnitine production can be rate-limiting for fatty acid metabolism (Fritz and McEwen 1959; Bremer 1983). In accordance with our previous results, we observed a decrease in steady-state levels of L-carnitine in Spen-KD|MYD compared to CTRL-KD|MYD (Figure 4A, Hazegh *et al.* 2017). Similarly, Spen-KD|HSD and Spen-KD|HFD had decreased levels of steady-state L-carnitine compared to matched-diet CTRL-KDs (Figure 4C-D). In contrast, HYD restored Spen-KD L-carnitine levels to those of matched CTRL-KD in the same diet (Figure 4B).

**Figure 4:**
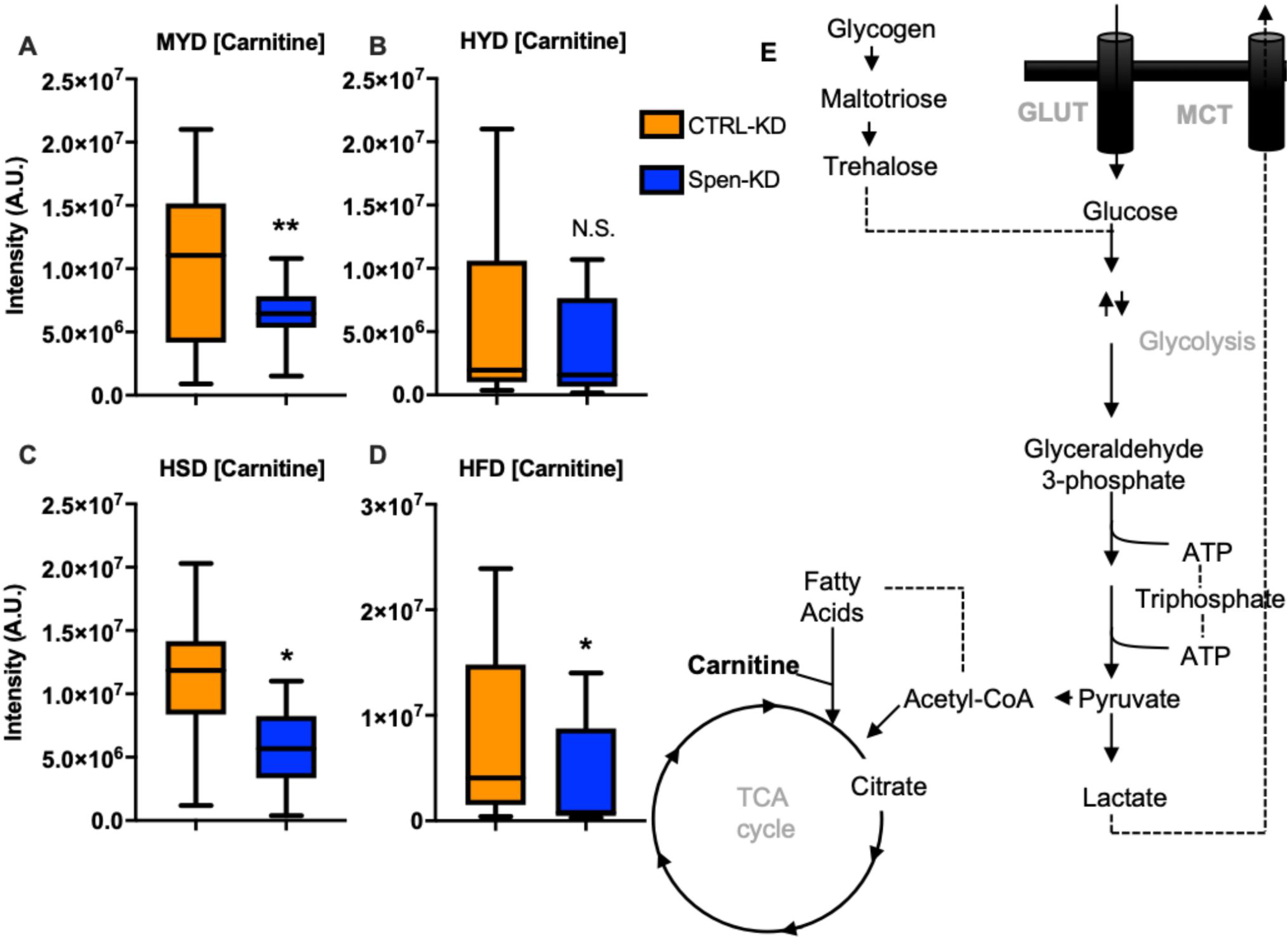
HYD rescues L-carnitine levels in Spen-KD. Levels of carnitine for in CTRL-KD (orange) and Spen-KD (blue) reared in A) Medium-Yeast Diet (MYD), B) High-Yeast Diet (HYD), C) High-Sugar Diet (HSD) and D) High-Fat Diet (HFD). Carnitine is an essential carrier for fatty acids into the mitochondrial matrix to facilitate β-oxidation of lipids as TCA cycle inputs. E) schematic summary of the role of carnitine in glycolysis. Student T-Test, *p<0.05, **p<0.01.

Additionally, metabolite set enrichment analysis (MSEA; MetaboAnalyist 4.0) showed in Spen-KD|MYD larvae an enrichment for pathways in amino acid metabolism and additional atypical metabolic processes (Figure 5A). Comparing the steady-state metabolome of Spen-KD|HYD to CTRL-KD|HYD, the metabolite set enrichment observed in the MYD was abolished and amino acid metabolism was no longer statistically significantly enriched in the Spen-KD|HYD (Figure 5B). Thus, amelioration of these metabolic steady-state defects could be the basis of the partial rescues we observed for development of Spen-KD raised on HYD. We also observed a large reduction in the metabolite steady-state variation between Spen-KD and CTRL-KD on the HSD and HFD (data not shown), suggesting that the supplemented quantity of sugar and fat within these diets is epistatic to the genetic influence of Spen depletion on metabolism.

**Figure 5:**
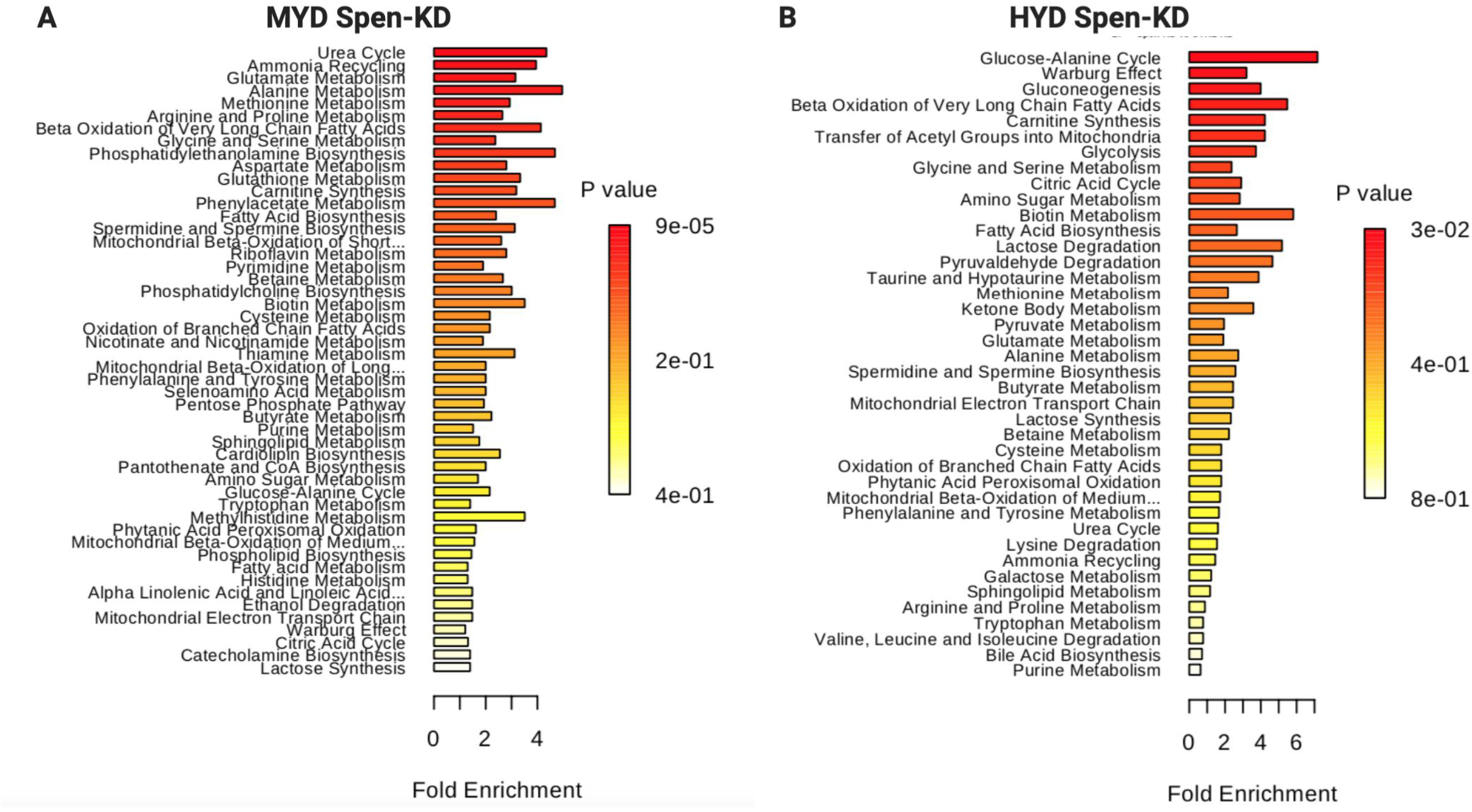
Diet x genotype-interactions dictate metabolic steady-state in third-instar larvae. Metabolite Set Enrichment Analysis (MSEA) Over Representation Analysis (ORA) generated by MetaboAnalyst 4.0. A) Spen-KD|MYD metabolite ORA compared to CTRL-KD|MYD B) and Spen-KD|HYD metabolite ORA compared to CTRL-KD|HYD.

To examine the global metabolomic differences within the same genotype between different diets we used orthogonal projections to latent structures data analysis (OPLS-DA; Figure 6). The metabolomic difference between Spen-KD on HYD and MYD was 15.4% (Figure 6C, F). Spen-KD on HSD was the least different from MYD with only 6.6% difference (Figure 6C, I). HFD drove the largest difference from MYD in Spen-KD at 24% difference (Figure 6C, L). CTRL-KD followed the same trend, with HSD being the least different (3.6%), followed by HYD (8.8%), and HFD was the most different (11.1%; Figure 6A). While CTRL-KD follows the same trend, the differences induced by diet challenge are much smaller than those observed in Spen-KD (e.g. CTRL-KD|HFD vs Spen-KD|HFD, 11.1% vs 24%). Therefore, the different diets induced similar patterns of metabolite changes regardless of genotype, but CTRL-KD was more resistant to change.

**Figure 6:**
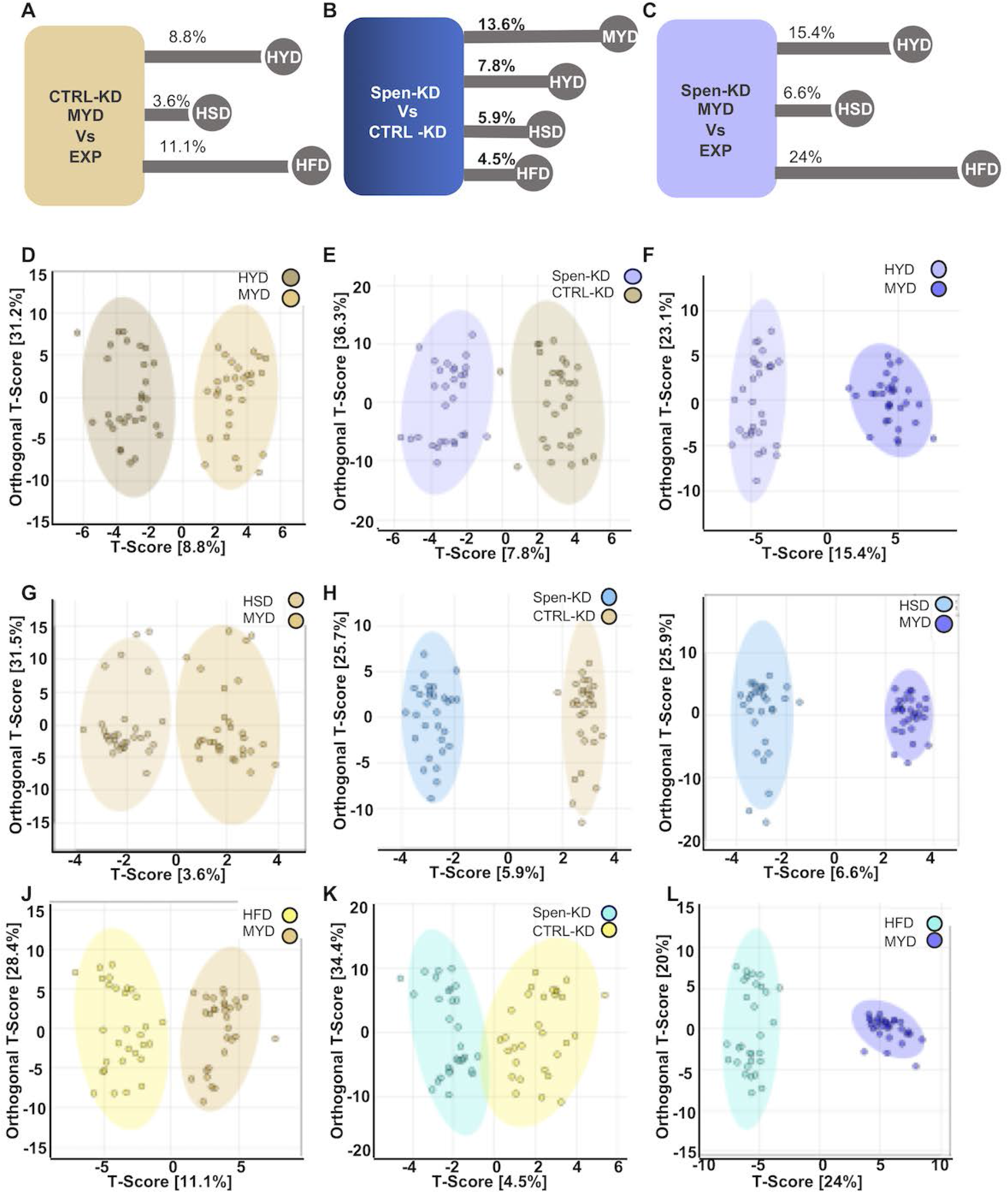
Orthogonal partial least squares discriminant analysis (OLPS-DA) comparisons of genotypes on diets and between genotypes. A) The metabolite differences (as percentage of all metabolites assayed) between CTRL-KD|MYD and CTRL-KD|EXP. (B-D) OLPS-DA of CTRL-KD|MYD compared to B) HYD C) HSD and D) HFD. E) Metabolite profile differences between Spen-KD and CTRL-KD on control MYD diet. (F-H) OLPS-DA of CTRL-KD (yellow) and Spen-KD (blue) on F) HYD G) HSD and H) HFD. I) Metabolite differences between Spen-KD|MYD and Spen-KD|EXP. (J-L) OLPS-DA of Spen--KD|MYD compared to Spen-KD| J) HYD K) HSD and L) HFD. T-score: variation between groups. Orthogonal T-Score: variation within the group.

We hypothesized that the developmental rescues observed reflected a rescue of the metabolic steady-state of Spen-KD. To this end, we compared diet-matched Spen-KD and CTRL-KD by OPLS-DA (Figure 6B). The largest difference between genotypes was on MYD (13.6%). HYD decreased the variability between Spen-KD and CTRL-KD to 7.8%, while HSD shrank it to 5.9% (Figure 6E, H). The least different was HFD at 4.5% (Figure 6K).

Altogether, our data suggest that diet challenge is epistatic to genotype influences on whole organism metabolism. CTRL-KD had more metabolic plasticity and was better able to resist metabolite change than Spen-KD. HFD, which drove severe developmental delays, provided the largest metabolic challenge to both genotypes and pushed them towards the most similar metabolic profiles (Figure 6B, K).

## DISCUSSION

Here we show that targeted gene–environment interactions can partially mitigate whole organismal pathophysiology. We show that supplementing the diet of Spen-KD individuals with macronutrients that fall within their metabolic capacities partially ameliorates the adiposity, development, and lifespan defects. We show that in Spen-KD animals a HYD partially restores diet-appropriate fat storage, and partially restores the timing of larval development at the wandering, pupariation, and eclosion time points. While HSD exacerbates the increased fat storage phenotype, it partially restores Spen-KD development during pupariation and eclosion and improves Spen-KD mated female lifespan. Diet challenge through macronutrient enrichment led to a decrease in metabolic steady-state differences between Spen-KD and CTRL-KD. The most pronounced differences were on MYD control-diet, which is optimized for larval development and adult fecundity in wildtype strains. HYD abolishes many of the metabolic steady-state differences between Spen-KD and CTRL-KD and brings the metabolic profile back towards more canonical larval pathways. Larvae upregulate metabolic pathways, using Warburg effect, to support their rapid development and accumulation of biomass (Tennessen *et al.* 2011). Spen-KD|HYD are enriched for metabolites involved in the Warburg effect, demonstrating a shift back to ‘canonical’ metabolic programming (Figure 5B).

We previously showed that Spen depletion in the fat body causes an obesity phenotype with defects in β-oxidation, increased protein catabolism, and increased glycolytic flux (Hazegh *et al.* 2017). These metabolic changes, most markedly the inability to properly liberate stored lipids, are accompanied by several developmental and lifespan defects. Here we demonstrate the successful application of targeted gene– environment interactions by using custom diets to alleviate Spen-derived developmental and lifespan defects.

We recapitulated our previous results and showed that Spen-KD|MYD had increased fat storage compared to CTRL-KD, as assayed by buoyancy (Figure 1A). As predicted, the Spen-KD|HSD and HFD groups were fatter than Spen-KD|MYD (Figure 1C-D, Table 1). Diet-induced obesity has been modeled in mammals through similar methods of supplementing diet with refined carbohydrates and fats (Levin *et al.* 1989; Buettner *et al.* 2007; Bortolin *et al.* 2018). These studies have shown that diets high in fat and carbohydrates are capable of producing obesity in rat and mouse models with similar pathology to human obesity and produce associated comorbidities and metabolic syndromes. In flies it has also been observed that carbohydrate- and fat-enriched diets induce obesity despite attenuated feeding behaviors, and drive metabolic syndromes and co-morbidities (Skorupa *et al.* 2008; Matzkin *et al.* 2011; Musselman *et al.* 2011; Birse *et al.* 2010). Our results are in agreement with these studies and further demonstrate that Spen-KD larvae follow these same expectations.

The CTRL-KD|HYD is fatter than the CTRL-KD|MYD, suggesting that the nutritionally enriched HYD is capable of inducing obesity (Figure 1B, Table 1). Conversely, Spen-KD|HYD larvae are less fat than Spen-KD|MYD, suggesting the HYD is protective against obesity in Spen-KD despite the nutrient enrichment (Figure 1B, Table 1). The difference between Spen-KD and CTRL-KD is smaller when reared on the HYD compared to MYD (Table 1), which suggests that the HYD decreases the difference in fat storage between Spen-KD and CTRL-KD. Therefore, HYD partially ameliorates the increased fat storage phenotype observed in Spen-KD larvae and narrows the adiposity differences between Spen-KD and CTRL-KD larvae.

Spen-KD|HSD is still significantly more fat than CTRL-KD|HSD, however carbohydrate enrichment also narrows the fat storage differences between Spen-KD and CTRL-KD (Figure 1C, Table 1). This was the smallest difference between diet-matched Spen-KD and CTRL-KD, suggesting that HSD-induced obesity originates from a different metabolic pathway. That is to say, HSD is epistatic to genotype in this investigation of gene-environment interaction. Supporting our observation, diet-supplemented carbohydrates have been used in mammalian and Drosophila models of diet-induced obesity (Surwit *et al.* 1995; Musselman *et al.* 2011; Nilsson *et al.* 2012).

HFD, as predicted, drives the largest increase in fat storage in both Spen-KD and CTRL-KD (Figure 1D, Table 1). This increase agrees with previous studies performed in mammalian and Drosophila models of diet-induced obesity, in which high-fat diets drive massive increases in fat storage in mice and rats (Levin *et al.* 1989; Buettner *et al.* 2007; Padmanabha and Baker 2014). Again, the Spen-KD|HFD store more fat than CTRL-KD|HFD, consistent with the trend observed in other diets.

Spen-KD displyed increased fat storage compared to CTRL-KD in all diet-matched assays (Figure 1). This result is consistent with our previously purposed model that Spen-KD generates a genetically underpinned obesity phenotype through dysregulated metabolism in fat body cells (Hazegh *et al.* 2017). As predicted, the HSD and the HFD induced significantly increased fat storage in both Spen-KD and CTRL-KD, consistent with the mammalian and Drosophila mode of diet induced obesity referenced above (Figure 1). This study demonstrates a gene-diet interaction between two genotypes and four diets. Most importantly, HYD partially rescues the increased fat storage phenotype in Spen-KD larvae. While Spen-KD|HYD still stores more fat than CTRL-MYD, HYD attenuates the development of genetic obesity and decreases the pathophysiological burden of excessive fat storage in these animals. The specificity of these gene-diet interactions provides support for the application of custom dietary intervention to treat obesity.

HYD and HSD partially rescue developmental delays associated with fat body-specific Spen depletion. To determine if macronutrient supplementation was capable of rescuing developmental defects in Spen-KD larvae, we measured developmental milestones in larvae reared on different diets. Spen-KD|HYD wander a median 17.4 hours earlier than Spen-KD|MYD. CTRL-KD|HYD wander a median 15.5 hours earlier than CTRL-KD|MYD. Taken together, this difference suggests that supplemented protein is broadly beneficial to developing larvae, regardless of genotype. The effect size takes into consideration the distribution of the experimental diet compared to the control diet developmental timing. HYD has a larger effect on CTRL-KD than Spen-KD, as denoted by the Cliff’s delta (d=-0.25). This result was surprising, as we initially hypothesized Spen-KD would experience greater benefits from a high protein diet than would CTRL-KD due to the upregulation of protein metabolism we previously observed (Hazegh *et al.* 2017). While protein catabolism can compensate in part for the decrease in available energy, it cannot fully rescue the metabolic defects observed in Spen-KD. Therefore, the beneficial effect of HYD on Spen-KD was smaller than CTRL-KD because it only addresses one part of organism-wide metabolic dysregulation.

HSD delays both Spen-KD (+2.8 hours from control) and CTRL-KD (+4 hour from control) wandering development (Figure 2B; Table 2). HSD negatively affected both genotypes, however it affected CTRL-KD more than Spen-KD (d=0.05). HSD drove the largest wandering delay in CTRL-KD, suggesting that excessive carbohydrates is the most detrimental macronutrient in early larval development in a control, or ‘wild-type’, genetic background.

As predicted, Spen-KD|HFD wandering was the most delayed from the MYD matched control (+5.2 hours from control). Since Spen-KD larvae cannot properly regulate lipid metabolism, it is not surprising that feeding larvae excess dietary fat has the strongest adverse effect on development (Figure 2C, Table 2, Hazegh *et al.* 2017). CTRL-KD|HFD is also delayed, however the effect size (d=-0.15) shows that HFD more strongly delays wandering in Spen-KD.

Essential amino acids derived from the diet (histidine, isoleucine, leucine, lysine, methionine, phenylalanine, threonine, tryptophan, valine, and arginine) must reach a threshold amount for larvae to commit to pupariation (Sang and King 1961). We hypothesize that the wandering development improvement measured in Spen-KD|HYD is a consequence of enriched essential amino acids. The wandering time of Spen-KD|HYD is similar to that of CTRL-KD|MYD, suggesting HYD improves early larval developmental timing to that of healthy controls.

It is likely that the HFD exacerbates the impaired lipid metabolic pathways within Spen-KD larvae, further exaggerating the developmental delays. Excess sugar is stored as a limited amount of glycogen and then shunted into lipogenesis to be stored as lipids (Acheson *et al.* 1982; Flatt 1995). We propose that the HSD exacerbates the obesity phenotype and lipid dysregulation through its contribution to increase *de novo* lipogenesis, but to a lesser degree than HFD because sugar can be shunted to fates other than TAGs (i.e. glycogen).

The only diets to significantly improve the pupariation and eclosion rates of Spen-KD larvae were HYD and HSD. While macronutrients are used as fuel for development, they also play an equally vital role as building blocks for biomass to support the 400-fold increase in body size the larvae undergo during their rapid development. We postulate that the increase in protein is insufficient to counteract the “starvation” effect caused by immobilized lipid stores, which are critical mediators of cell growth and division by providing phospholipid, glycolipid, and glycosphingolipid precursors. Glycosphingolipids are necessary for proper cell cytokinesis in mammals, and studies in Drosophila have shown proper neuronal expansion and remodeling requires the lipid metabolism pathway SREBP (Ziegler *et al.* 2017; Huang *et al.* 2018). We postulate that the persisting developmental delays are consequence of dysregulated lipid metabolism, and while HYD and HSD can improve development, it cannot compensate for this critical failure.

HYD partially rescued the Spen-KD excessive fat storage phenotype, and partially rescued delayed pupariation and eclosion, however it became detrimental to the longevity of both male and mated female Spen-KD adults (Figure 3A, D, Table 3). This is in accordance with previous data suggesting a high protein diet has negative longitudinal impacts on lifespan (Kitada *et al.* 2019). Conversely, CTRL-KD mated female and male lifespans are unaffected by HYD (Figure 3A, D, Table 3). This result suggests a sex-specific sex–diet–genotype interaction. We hypothesize that this sex difference in CTRL-KD lifespan is due to the unique metabolic needs of egg production and egg laying sustained by mated females as previous studies have shown that dietary amino acids are crucial to these processes (Lee *et al.* 2008; Haussmann *et al.* 2013). We hypothesize that genetic background, food media preparation, and assay conditions contribute to the heterogeneity observed in diet lifespan studies further emphasizing the importance of controlling gene-diet-environment-interactions in experimental designs.

HFD is extremely deleterious to lifespan of both sexes and genotypes of flies assayed (Driver and Cosopodiotis 1979; Woodcock *et al.* 2015). Our data agrees with precedent in the literature, with severely attenuated lifespans in both males and mated females of both genotypes. Males survived better on HFD than genotype-matched mated females. Previous studies showed major transcriptome differences between males and mated females reared on HFD: females upregulated fatty acid metabolism and their stress response while males downregulated stress response genes (Heinrichsen *et al.* 2014; Stobdan *et al.* 2019). Our data suggests that Spen-KD flies maintain this sex-specific sensitivity to dietary fat intake.

### Diet composition is a primary variable in the metabolome at steady-state

By challenging Spen-KD and CTRL-KD larvae with HSD, HFD, and HYD we were largely able to abolish the influence of genetics on the metabolic steady-state of the animals and achieved strikingly similar metabolomic profiles between the two genotypes. The most different metabolite profiles occur on MYD, with 13.6% of metabolites being expressed differently between Spen-KD and CTRL-KD (Figure 6B).

Dietary extremes appear epistatic to more nuanced genetic regulators of metabolism, such as Spen-KD. Larvae reared on the HFD are least different between Spen-KD and CTRL-KD genotypes, differing in only 2 of 140 metabolites assessed (acetylcholine and L-Carnitine) with a global 4.5% difference between genotypes. This is an interesting observation when paired with the observation that Spen-KD|HFD are significantly more fat than CTRL-KD|HFD controls. We postulate that Spen-KD is less resilient to diet challenge and develops more severe pathophysiological obesity phenotypes due to their defective lipid mobilization.

The role of Spen in metabolism changes as metabolic needs change through maturation. HYD partially rescues pupation and eclosion timing of Spen-KD larvae but a significant negative impact on adult lifespan, which is in agreement with existing literature on the influence of protein in the Drosophila lifecycle (Lee *et al.* 2008; Kitada *et al.* 2019). Spen-KD is more sensitive to the beneficial developmental effects of HYD than CTRL-KD in whole organismal energy homeostasis, pupariation, and eclosion.

HSD improves pupariation and eclosion larval development as well as partially rescues Spen-KD adult lifespans. Unlike the high-protein diet, excess carbohydrates is beneficial and confers starvation resistance in mated females, and to a lesser extent males (Chandegra *et al.* 2017). As described in our previous study, Spen-KD alters carbohydrate metabolism and changes glycolytic flux (Hazegh *et al.* 2017). We see in the adult stage wherein HSD is a more favorable diet composition for longevity of Spen-KD mated females than do CTRL-KD. This is not conserved in males, which is not entirely surprising given the significantly smaller benefit of increased sugar conferred to adult males in general (Chandegra *et al.* 2017). HSD alters the metabolite profiles of Spen-KD and CTRL-KD the least, however Spen-KD|HSD changes twice as much CTRL-KD|HSD compared to MYD matched controls (Figure 6A, C).

While we are interested in genetically-driven forms of obesity, we seek to understand the functions of these genes in a whole-organism metabolic context in order to leverage this molecular understanding towards designing diet interventions that ameliorate pathologies. This study demonstrates the feasibility of gene-macronutrient interventions as a treatment for obesity and related co-morbidities resulting from a specific genetic perturbation.

## ACKNOWLEDGEMENTS

We would like to thank the Bloomington Stock Center, for providing fly stocks. Michael McMurray for his helpful comments on the manuscript. This work was supported by NIH-T32-GM008730 and the Victor W. and Earleen D. Bolie Graduate Scholarship to K.E.H., NIH-T32-GM008730 and RNA Scholar of the RNA Bioscience Initiative, University of Colorado School of Medicine to C.M.G. A.D. was supported by funds from the Boettcher Webb-Waring Investigator Award, NIH/NIGMS Grant RM1GM131968 and NIH/NHLBI Grant R01HL146442. J.M.T. is supported by the RNA Bioscience Initiative at the University of Colorado Anschutz Medical Campus and a Webb-Waring Early Career Investigator Award from the Boettcher Foundation (AWD-182937). T.R. is supported by NIH/NIDDK Grant R01DK106177 and a pilot grant from the RNA Bioscience Initiative, University of Colorado School of Medicine.

